# An arthropod-specific TMEM16 protein accelerates olfactory response termination in *Drosophila*

**DOI:** 10.64898/2026.05.21.727025

**Authors:** Ying Lei, Jing Lei, Tianbang Li, Makoto Tominaga, Wynand Van Der Goes Van Naters, Tatsuhiko Kadowaki

**Author notes:** Correspondence: Wynand Van Der Goes Van Naters, School of Biosciences, Cardiff University, Sir Martin Evans Building, Museum Avenue, Cardiff, CF10 3AX, UK, Tatsuhiko Kadowaki, Department of Biosciences and Bioinformatics, School of Science, Xi’an Jiaotong-Liverpool University, 111 Ren’ai Road, Suzhou Dushu Lake Higher Education Town, Jiangsu Province 215123, China. Academy of Pharmacy, Xi’an-Jiaotong Liverpool University, Suzhou Dushu Lake Higher Education Town, Jiangsu Province, 215123, China (Tianbang Li). Thermal Biology Research Group, Nagoya Advanced Research and Development Center, Nagoya City University, Kawasumi 1, Mizuho-cho, Mizuho-ku, Nagoya 467-8601, Japan (Makoto Tominaga).

## Abstract

Rapid termination of odor responses is essential for accurate olfactory coding in insects, where odorant receptors function as ligand-gated ion channels rather than G protein-coupled receptors. However, the mechanisms that restore olfactory receptor neuron (ORN) excitability after stimulation remain poorly understood. Here, we identify DmTMEM16O (CG6938), an arthropod-specific member of the TMEM16 (Anoctamin) family, as a key regulator of ORN response termination in *Drosophila melanogaster*. DmTMEM16O is highly enriched in Orco-positive ORNs in both larval and adult olfactory organs. Loss of DmTMEM16O prolongs odor-evoked neuronal activity, increasing decay time constants and causing persistent depolarization. DmTMEM16O mutant ORNs fail to resolve repeated odor stimulation and show impaired temporal coding, accompanied by reduced behavioral responses to both attractive and aversive odorants. Cell-specific rescue demonstrates that DmTMEM16O acts within ORNs to accelerate response termination. Although DmTMEM16O does not exhibit detectable Ca^2+^-activated chloride channel activity in heterologous cells, our results support a model in which it increases membrane conductance during the decay phase of the response, thereby shortening the membrane time constant and promoting rapid repolarization. This function is consistent with a role for chloride influx in insect ORNs, in contrast to mammalian systems where TMEM16B-mediated chloride efflux amplifies depolarization. Together, our findings identify DmTMEM16O as a lineage-specific regulator of ORN dynamics that enables precise temporal coding in insect olfaction.

## Introduction

Olfaction is essential for insect survival and reproduction, guiding behaviors such as food seeking, mate recognition, oviposition, and avoidance of predators and toxins. In *Drosophila melanogaster*, odorants are detected by olfactory receptor neurons (ORNs) located in the antennae and maxillary palps of adults and in the dorsal organ ganglia (DOG) of larvae. These neurons encode odor identity, intensity, and temporal structure and relay this information to higher-order circuits in the antennal lobe. Because natural odor signals are highly dynamic, precise control of both activation and termination of ORN responses is required to preserve temporal fidelity and ensure accurate behavioral outputs [1, 2].

Insect olfactory transduction differs fundamentally from that of mammals. Mammalian odorant receptors are G protein-coupled receptors (GPCRs) that initiate second messenger signaling cascades, whereas insect odorant receptors (ORs) function as heteromeric ligand-gated ion channels composed of a tuning receptor and the conserved co-receptor Orco. Odor binding directly opens these channels, allowing rapid cation influx and membrane depolarization [3–5]. While the mechanisms of activation are well established, the processes that govern response termination and membrane repolarization in insect ORNs remain incompletely understood. In particular, how ORNs rapidly reset after odor offset to maintain temporal resolution is a central unresolved question.

The TMEM16 (Anoctamin) family comprises membrane proteins that function as calcium-activated chloride channels (CaCCs) or phospholipid scramblases [6]. In vertebrates, TMEM16A and TMEM16B are well-characterized CaCCs with important roles in epithelial physiology and sensory systems, including olfaction [7, 8]. In mammalian ORNs, odor stimulation activates a cAMP-dependent signaling cascade that leads to Ca^2+^ influx through cyclic nucleotide–gated channels. This Ca^2+^ signal in turn activates TMEM16B [9, 10]. As the mammalian ORNs’ ciliary Cl^−^ concentration is high [11], opening of TMEM16B results in Cl⁻ efflux, which further depolarizes the neuron and amplifies the odor response [12]. Thus, TMEM16B is required to sustain and enhance ORN activation. In *Drosophila*, conserved TMEM16A/B-like protein, Subdued, has been implicated in ion transport and membrane physiology, exhibiting both ion channel and lipid scramblase activities [13–15]. However, whether TMEM16 family members contribute to olfactory signaling remains unclear.

Among TMEM16 family members, TMEM16O represents a distinct arthropod-specific clade that is absent from vertebrates and other non-arthropod lineages [16]. This restricted evolutionary distribution suggests that TMEM16O emerged to support physiological processes unique to arthropods. One such process in insects is olfactory transduction mediated by ionotropic OR–Orco channels, which imposes distinct demands on membrane repolarization and temporal signal resolution. A conductance that accelerates repolarization would effectively reduce the membrane time constant and enable rapid recovery of ORNs following stimulation, thereby preserving temporal coding in fluctuating odor environments.

Here, we characterized the function of the *Drosophila melanogaster* TMEM16O homolog, CG6938 (*DmTMEM16O*). We show that *DmTMEM16O* is highly enriched in Orco-positive ORNs of both larval and adult olfactory organs and is required for rapid termination of odor-evoked responses. Loss of *DmTMEM16O* prolongs neuronal activity, increases decay time constants, and impairs temporal resolution and olfactory behavior. Although DmTMEM16O does not exhibit canonical Ca^2+^-activated chloride channel activity in a heterologous system, our findings support a model in which it contributes to membrane conductance during response decay, thereby accelerating repolarization. These results identify DmTMEM16O as a functionally specialized, arthropod-specific regulator of ORN dynamics that supports the unique physiological demands of insect olfactory signaling.

## Results

### DmTMEM16O-Gal4 is expressed in olfactory receptor neurons (ORNs) of fruit fly larva and adult

To determine where *DmTMEM16O* (CG6938) is expressed in *D. melanogaster*, we first constructed a *DmTMEM16O-Gal4* stock, in which Gal4 is driven by the 5′ upstream region (2209 bp) of *DmTMEM16O*. By crossing this stock with a 20×*UAS*-6×*GFP* stock [17], we found that *DmTMEM16O-Gal4* is highly expressed in the dorsal organ ganglia (DOG) of 1st instar larvae (Fig. 1A). Since the larval DOG of *D. melanogaster* contains both ORNs and ionotropic receptor (IR)-expressing sensory neurons [18, 19], we tested whether *DmTMEM16O* colocalizes with either olfactory receptor (OR)-positive or IR-positive neurons. IR-positive neurons were identified using tdTomato driven by the 5′ upstream region (1422 bp) of *IR25a* (IR25a-tdTomato). As shown in Fig. 1B and C, DmTMEM16O is present in Orco-positive ORNs but not in IR25a-positive sensory neurons. In addition to ORNs in the DOG, DmTMEM16O was also detected in pharyngeal neurons (Fig. 1D).

**Figure 1.**
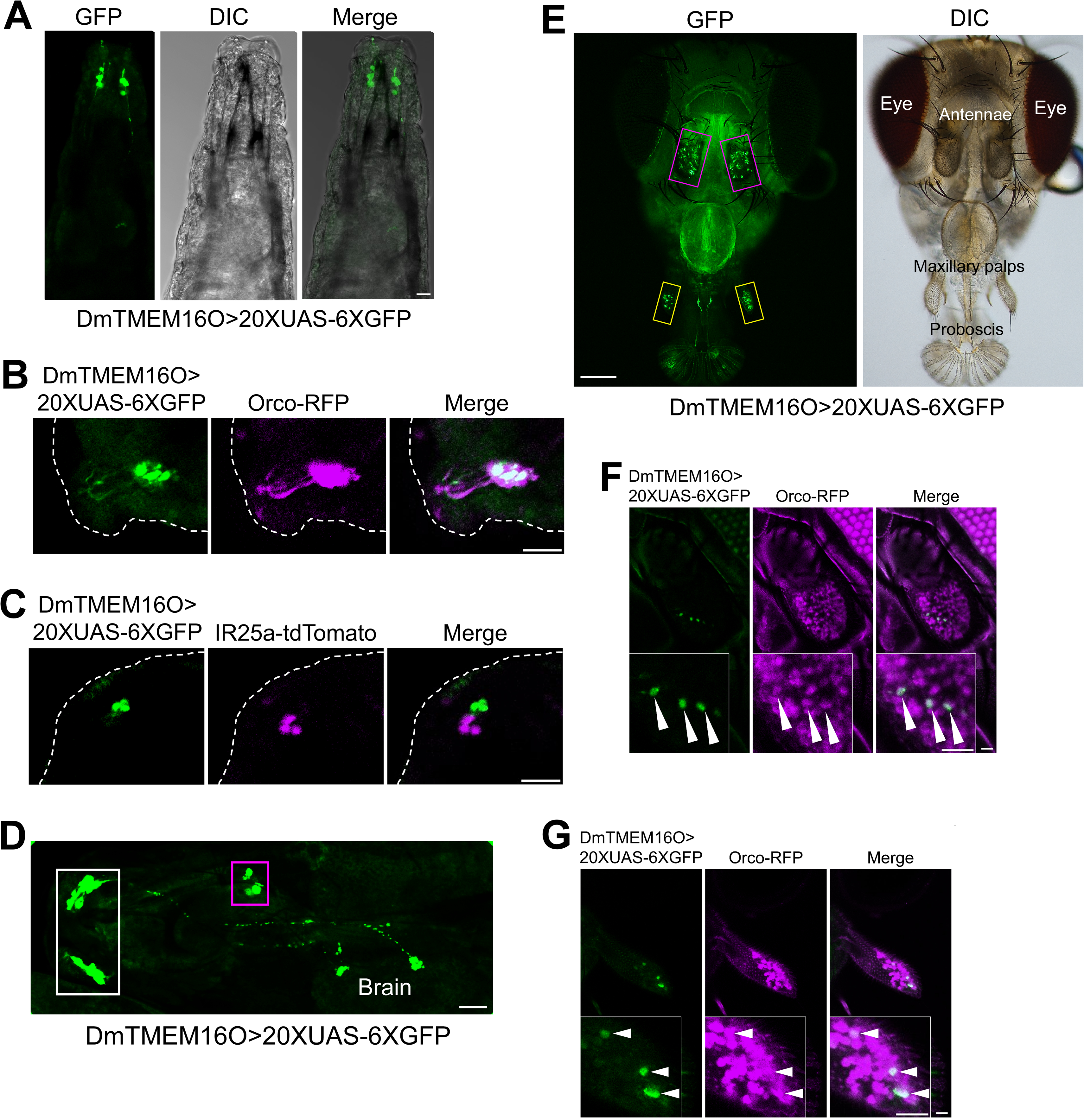
DmTMEM16O is expressed in Orco-positive olfactory receptor neurons. Larval expression is shown in panels A–D and adult expression in panels E–G. (A) Expression of DmTMEM16O in the dorsal organ ganglia (DOG) of first instar larvae visualized using DmTMEM16O-Gal4 driving 20×UAS-6×GFP. Differential interference contrast (DIC) and merged images are also shown. Anterior is at the top. (B and C) DmTMEM16O-Gal4-driven GFP signal co-localizes with Orco-positive olfactory receptor neurons (ORNs; Orco-RFP), but not with IR25a-positive neurons (IR25a-tdTomato), in the larval DOG (Merge). The outline of the larva is indicated by a white dotted line. (D) DmTMEM16O-Gal4-driven GFP signal is also detected in pharyngeal gustatory receptor neurons (purple square). ORNs in the DOG (white square) project axons to the antennal lobes of the brain. Anterior is at the left. (E) DmTMEM16O-Gal4-driven GFP expression in adult olfactory organs, including the antennae (purple squares) and maxillary palps (yellow squares). A DIC image showing head structures is also included. DmTMEM16O-Gal4-driven GFP co-localizes with Orco-positive ORNs in the antenna (F) and maxillary palp (G) (Merge). Insets show higher magnification views of DmTMEM16O-positive neurons (white arrowheads). Scale bars, 20 μm except 100 μm in (E).

DmTMEM16O is expressed in adult antennae and maxillary palps (Fig. 1E). As in larvae, DmTMEM16O is present in Orco-positive ORNs (Fig. 1F and G) in both antennae and maxillary palps. Thus, DmTMEM16O is enriched in ORNs of both larval and adult fruit flies. These results are consistent with Fly Cell Atlas data showing that *DmTMEM16O* mRNA is highly expressed in various ORNs of the adult fly antenna [20].

### Loss of DmTMEM16O impairs the termination of ORN responses to odors

Given that DmTMEM16O is highly expressed in ORNs, it may be associated with olfactory behavior in fruit flies. To investigate its physiological role, we generated a DmTMEM16O mutant fly in which the coding region corresponding to amino acids 153–1016 was deleted using CRISPR by expressing two gRNAs targeting exon 3 and exon 10. The deleted region contains the Anoctamin dimerization domain and most of the Anoctamin family domain, suggesting that this deletion represents a null allele (Supplementary Figure 1). Homozygous mutant flies (*DmTMEM16O^Δ153–1016^*) were viable and fertile.

We measured the responses of large basiconic sensilla (ab1, ab2, and ab3) in the antenna to five odors in wild-type (WT, *Oregon-R*) and *DmTMEM16O^Δ153–1016^* flies (backcrossed to *Oregon-R* five times) using single sensillum recording (SSR). Ab1 sensilla hold four neurons while ab2 and ab3 each house a pair of neurons, and the odors tested are strong ligands for some neurons in these sensilla [21, 22] (Supplementary Figure 2). In the responses of ab1 sensilla (ab1D neuron) to methyl salicylate, ab2 sensilla (ab2A neuron) to ethyl acetate, ab2 sensilla (ab2B neuron) to ethyl 3-hydroxybutyrate, ab3 sensilla (ab3A neuron) to ethyl butyrate, and ab3 sensilla (ab3B neuron) to 2-heptanone, both spiking activity and slow voltage deflection decayed more slowly after odor offset in the mutant than in WT flies (Fig. 2A). A similar phenotype was also observed in thin basiconic sensilla of the maxillary palp. The responses of pb1 sensilla to racemic 3-octanol (activating pb1A neuron) and 4-methyl phenol (activating pb1B neuron) terminated more slowly in the mutant flies (Fig. 2B). Accordingly, the exponential decay constants of the responses of basiconic sensilla to all of these compounds were longer in the mutant flies (Fig. 2C and D). These results demonstrate that DmTMEM16O normally accelerates the shutoff of ORN responses to odors.

**Figure 2.**
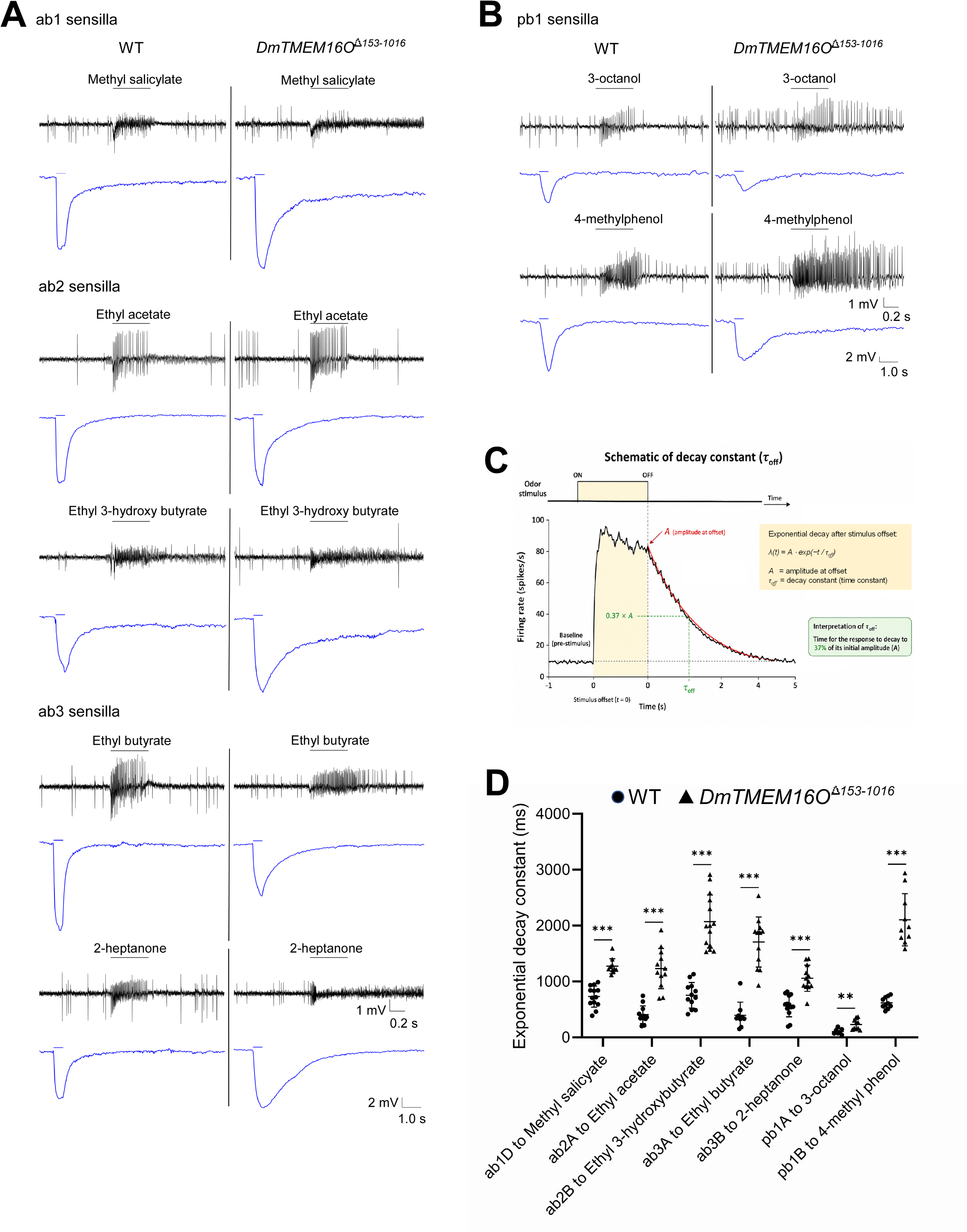
DmTMEM16O is required for rapid termination of odor-evoked responses (A) Representative single sensillum recording (SSR) traces from ab1, ab2, and ab3 sensilla in response to the indicated odorants. In *DmTMEM16O^Δ153–1016^* flies, both spiking activity (black) and receptor potentials (blue) decay more slowly after odor offset compared with WT. (B) Representative SSR traces from pb1 sensilla in the maxillary palp in response to racemic 3-octanol and 4-methyl phenol, showing similarly prolonged response termination in *DmTMEM16O^Δ153–1016^* flies. (C) Schematic representation of the measurement of the exponential decay constant (*τ_off_*). (D) Comparison of *τ_off_* values for ORNs in large and thin basiconic sensilla in response to the indicated odorants, demonstrating significantly prolonged response decay in *DmTMEM16O^Δ153–1016^* flies. Error bars represent mean ± SD (n = 9–14). \*\**P* < 0.005 and \*\*\**P* < 0.001 (Mann–Whitney U test).

We next tested whether the SSR phenotypes of *DmTMEM16O^Δ153–1016^* could be rescued by expressing DmTMEM16O specifically in ab3A neurons using *OR22a-Gal4*. In the parental lines (*UAS-DmTMEM16O* and *OR22a-Gal4* in the *DmTMEM16O^Δ153–1016^*background), responses of ab3 sensilla to both ethyl butyrate and 2-heptanone showed slow termination, similar to that observed in the mutant (Fig. 2A). In contrast, in the progeny expressing DmTMEM16O in ab3A neurons, the response to ethyl butyrate - but not to 2-heptanone which acts on the ab3B neuron - terminated more rapidly (Fig. 3A). Consistent with this, the exponential decay constant of ab3A responses to ethyl butyrate, but not ab3B responses to 2-heptanone, was significantly reduced in the rescued flies compared with the parental lines (Fig. 3B). These results demonstrate that DmTMEM16O specifically rescues the response kinetics of ab3A neurons (Or22a-positive), but not ab3B neurons (Or22a-negative).

**Figure 3.**
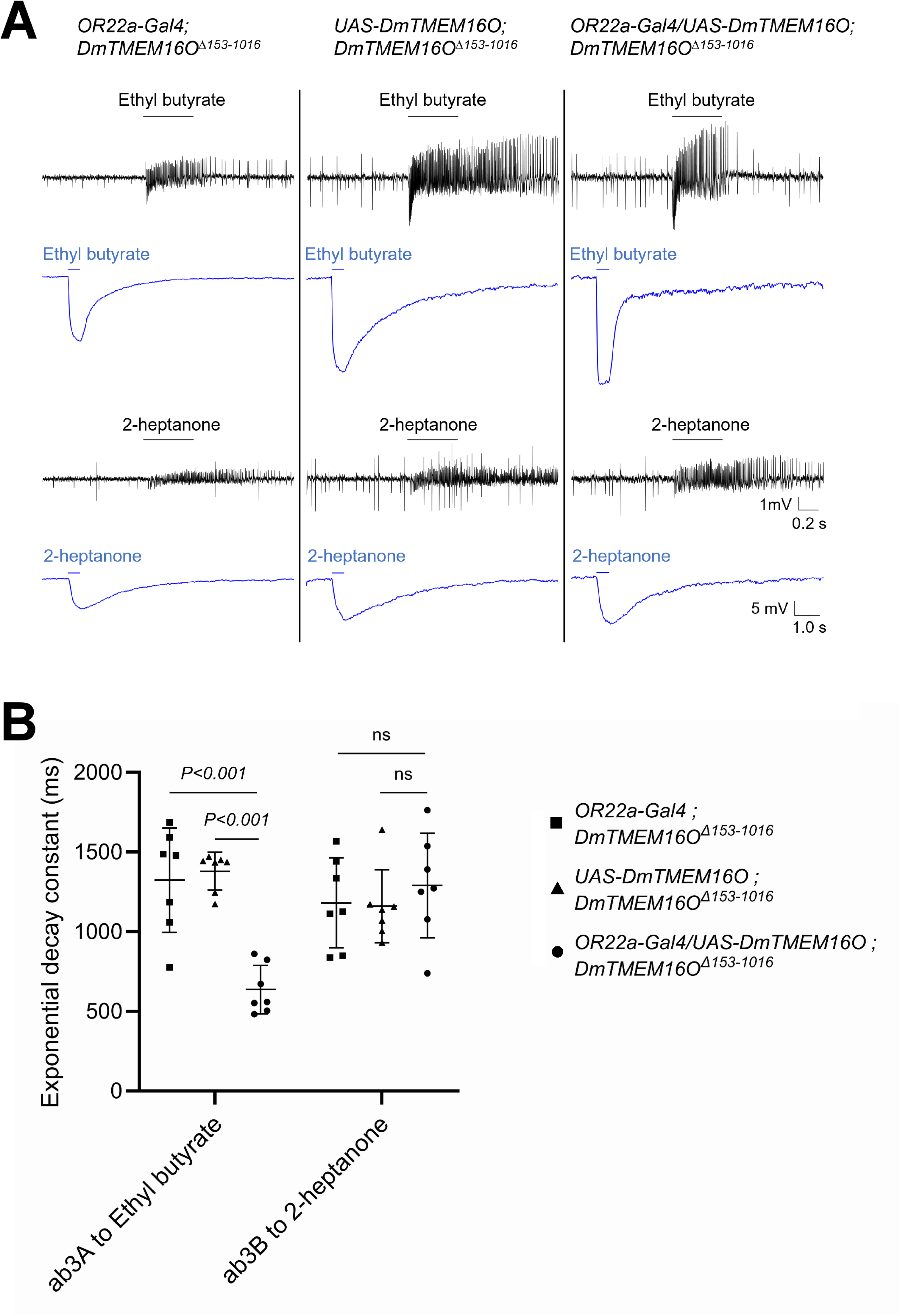
Cell-autonomous rescue of response termination in ab3A neurons. (A) SSR traces (black) together with receptor potentials (blue) from ab3 sensilla in response to ethyl butyrate and 2-heptanone. Parental lines (*UAS-DmTMEM16O* and *OR22a-Gal4* in the *DmTMEM16O^Δ153–1016^* background) exhibit slow response termination. Expression of DmTMEM16O under control of *OR22a-Gal4* restores rapid termination of responses in ab3A neurons to ethyl butyrate, but not in ab3B neurons to 2-heptanone; note *OR22a-Gal4* expresses only in ab3A. (B) Comparison of exponential decay constants. Decay of ab3A responses to ethyl butyrate is significantly faster in rescued flies compared to parental controls (*P* < 0.001), whereas ab3B responses to 2-heptanone are unchanged (ns: not significant) (Mann-Whitney U test). Error bars represent mean ± SD (n = 7).

Since DmTMEM16O mutant ORNs remained activated during the off-time of odor stimulation, we further examined how WT and mutant ORNs respond to five repeated odor stimuli. WT ab3A neurons resolved repeated temporal stimulation with ethyl butyrate well. In contrast, mutant ab3A neurons continued firing throughout the entire stimulation period and failed to resolve temporal odor stimulation (Fig. 4A-C).

**Figure 4.**
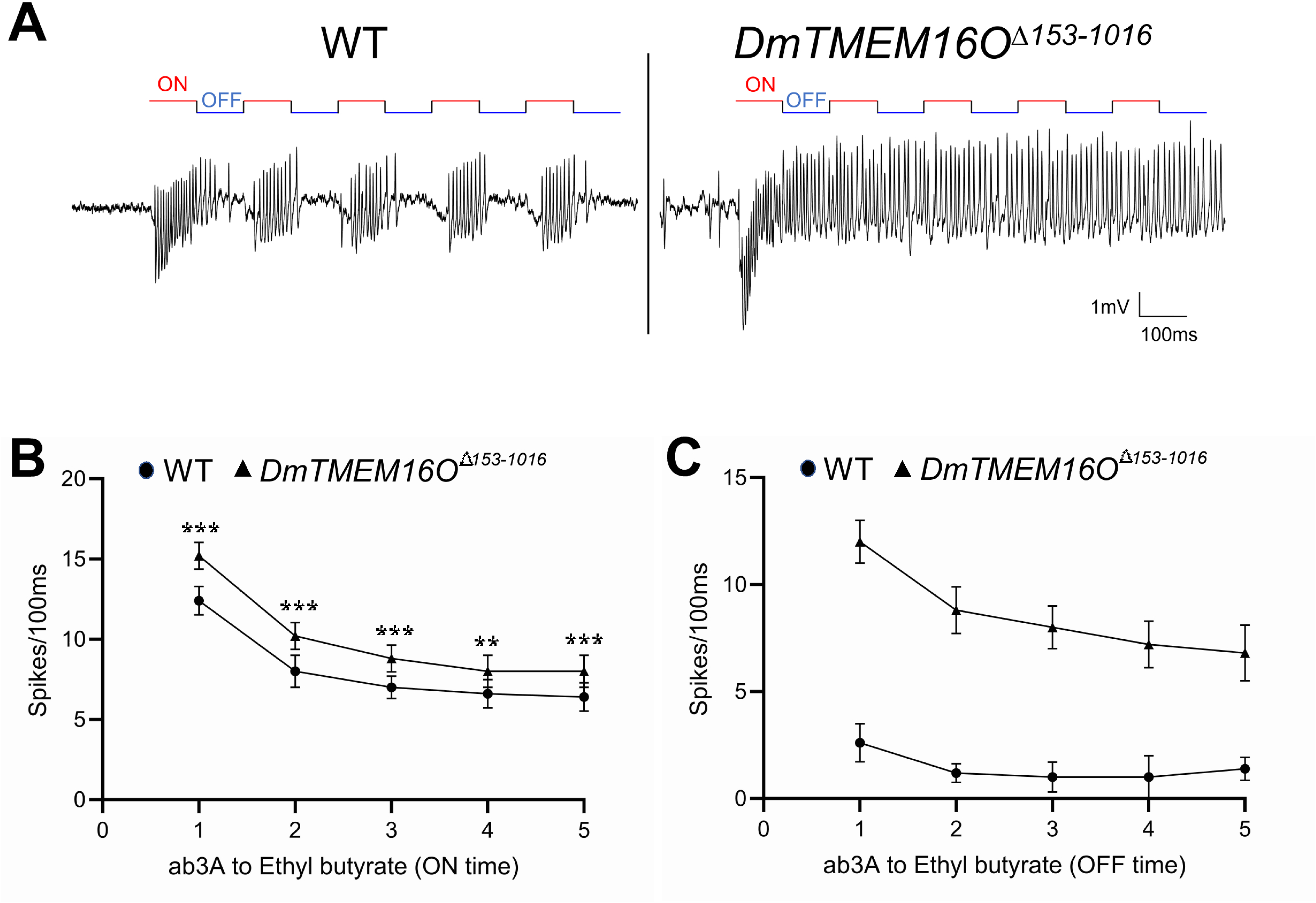
DmTMEM16O is required for temporal resolution of odor stimuli. (A) SSR recordings from ab3 sensilla during five repeated stimuli with ethyl butyrate (100 msec each for ON and OFF). WT neurons resolve individual pulses and return to baseline between stimuli. In *DmTMEM16O^Δ153–1016^*flies, ORNs exhibit sustained firing, failing to return to baseline between pulses. The number of spikes during the five ON (B) and OFF (C) times for WT and *DmTMEM16O^Δ153–1016^* flies. Error bars represent mean ± SD (n = 5). \*\**P* < 0.003 and \*\*\**P* < 0.001 (Brunner–Munzel test).

### *DmTMEM16O^Δ153–1016^* flies show reduced behavioral responses to odors

We next asked whether the altered ORN response phenotype observed in *DmTMEM16O^Δ153–1016^* flies by SSR affects olfactory behavior. Behavioral responses of adult flies to specific odors were examined using a standard T-maze assay. Adult flies were attracted to the ester compounds ethyl acetate, ethyl butyrate, and ethyl 3-hydroxybutyrate, as well as to 2-heptanone, but avoided racemic 3-octanol, methyl salicylate, and benzaldehyde (Fig. 5). The behavioral responses of *DmTMEM16O^Δ153–1016^*flies to these odors were significantly weaker than those of WT flies (Fig. 5), except at the low concentration of racemic 3-octanol (10^-4^ dose).

**Figure 5.**
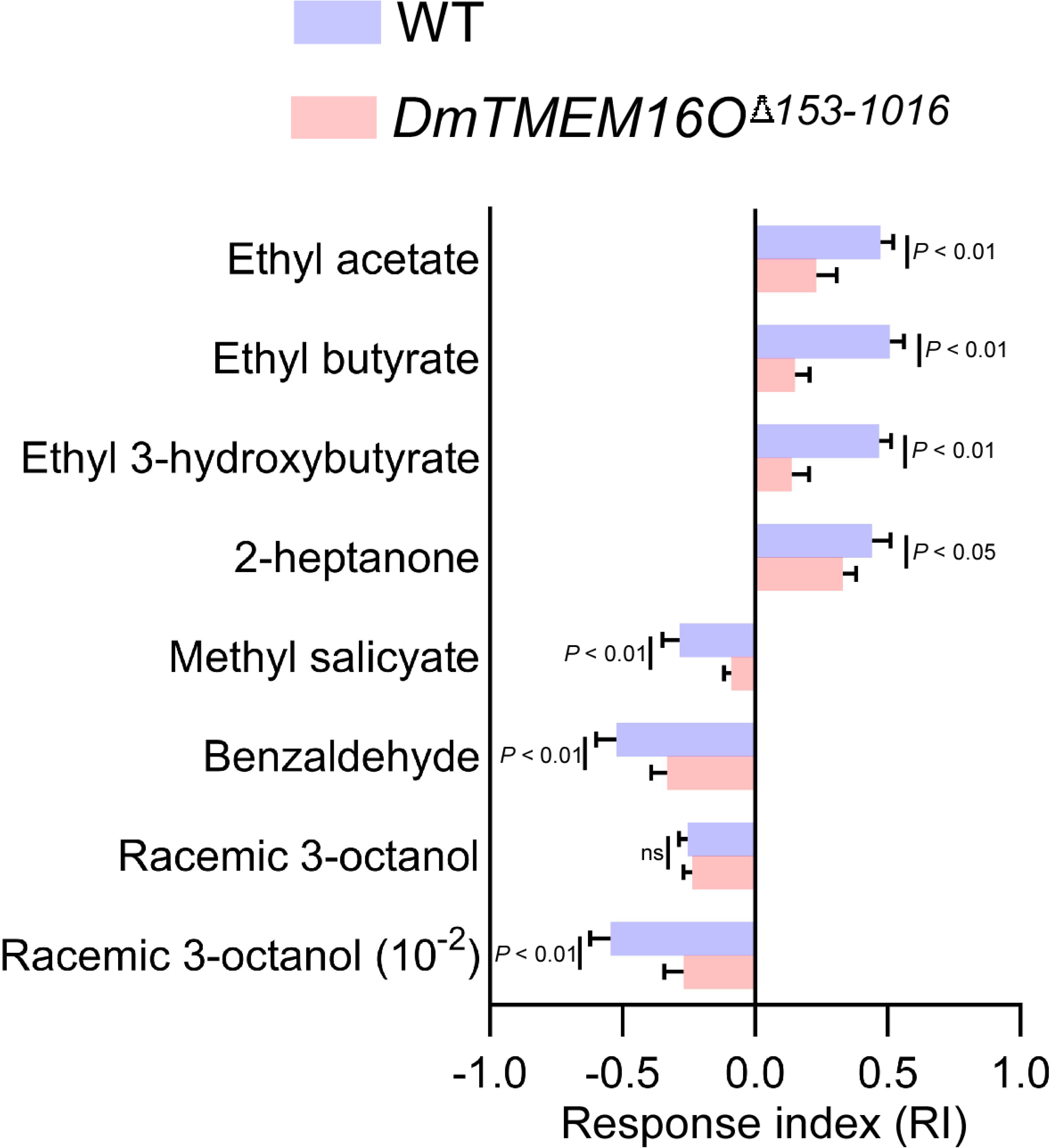
DmTMEM16O is required for normal olfactory behavior. Behavioral responses measured by T-maze assay (Response index). WT flies are attracted to fruit-related odorants (ethyl acetate, ethyl butyrate, ethyl 3-hydroxybutyrate, 2-heptanone) and avoid some other odorants (racemic 3-octanol, methyl salicylate, benzaldehyde). *DmTMEM16O^Δ153–1016^* flies show significantly reduced responses to these odorants, except at low concentrations of racemic 3-octanol (ns at 10⁻⁴). Error bars represent mean ± SD (n = 5). *P-*values by Mann-Whitney U test are also indicated.

### In a heterologous expression system DmTMEM16O does not act as a calcium-dependent chloride channel (CaCC), although a different DmTMEM16 paralog does

Based on the above results, DmTMEM16O may function as a CaCC that speeds repolarization of ORNs after odor activation. We therefore tested whether DmTMEM16O functions as a CaCC using whole cell patch-clamp analysis in a heterologous expression system in HEK293T cells. As shown in Figure 6A, DmTMEM16O did not generate any detectable current in the presence of 100 μM intracellular Ca^2+^ under various voltage conditions.

**Figure 6.**
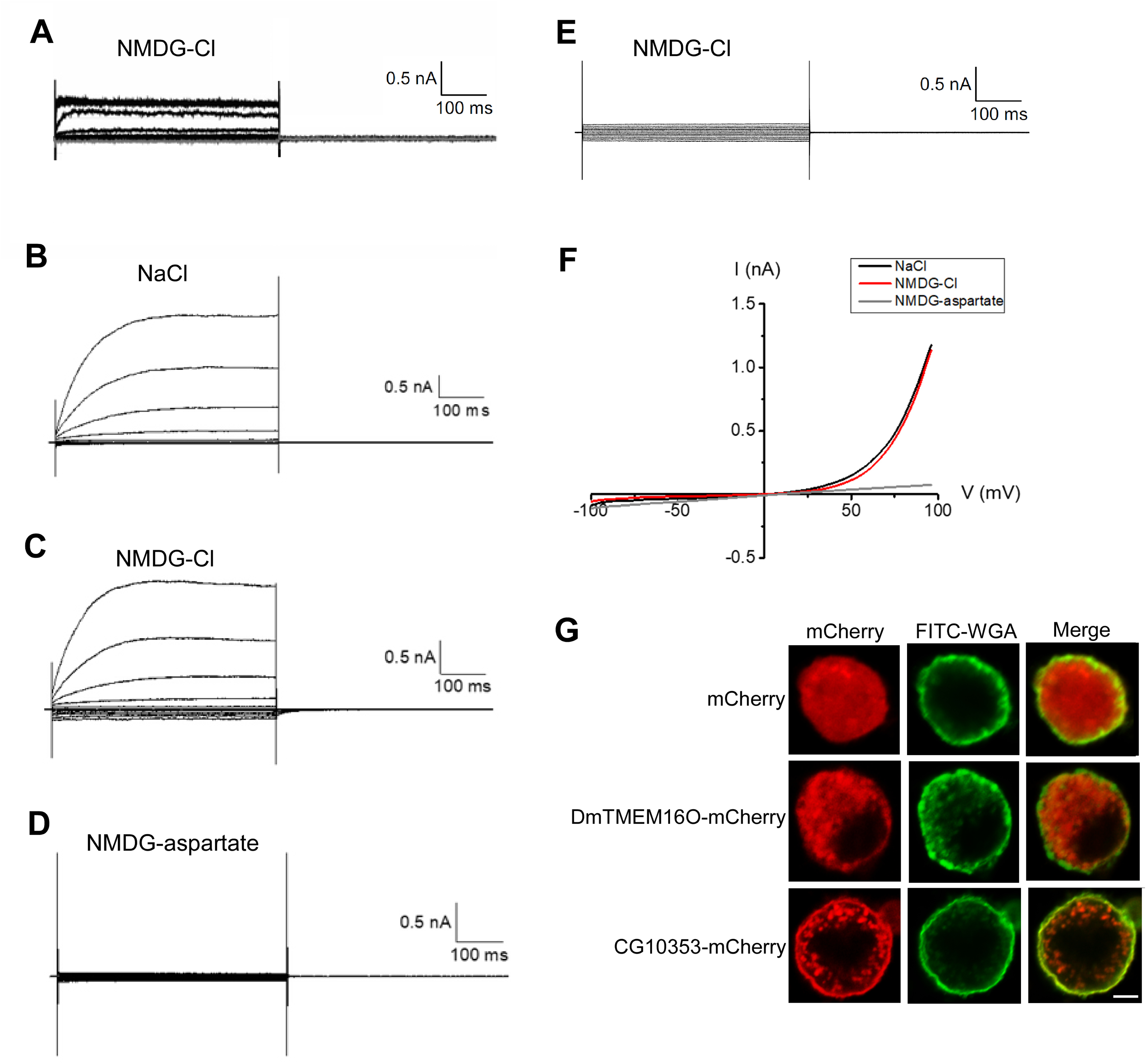
CG10353, but not DmTMEM16O, is a calcium-dependent chloride channel in HEK293T cells. (A) Whole-cell recordings from HEK293T cells expressing DmTMEM16O. No detectable currents are observed in the presence of 100 μM intracellular Ca²⁺ during voltage steps from −100 to +100 mV. Whole-cell recordings from cells expressing CG10353 show robust currents in Cl⁻-containing extracellular solutions with either NaCl (B) or NMDG-Cl (C) but not when Cl⁻ is replaced with aspartate, NMDG-aspartate (D). All recordings were conducted in the presence of 100 μM intracellular Ca²⁺. (E) CG10353-mediated currents are abolished in the absence of intracellular Ca^2+^. (F) Current–voltage relationship of CG10353 currents shows an outwardly rectifying Cl⁻ conductance. (G) Subcellular localization of mCherry-tagged proteins. CG10353-mCherry localizes to the plasma membrane as indicated by co-localization with FITC-WGA, whereas mCherry and DmTMEM16O-mCherry do not (Merge). Scale bar, 5 μm.

As a reference, we also examined CG10353, one of the two *D. melanogaster* TMEM16A/B family members [16]. Another family member, Subdued, was previously shown to possess both lipid scramblase and ion channel activities [15]. CG10353 generated currents in the presence of NaCl or NMDG-Cl, but not NMDG-aspartate, under voltage-clamp conditions from −100 to +100 mV (Fig. 6B–D). The current observed with NMDG-Cl was absent in the absence of intracellular Ca^2+^ (Fig. 6E). The I–V relationship showed a strongly outwardly rectifying Cl⁻ conductance that was abolished when extracellular and intracellular Cl⁻ was replaced with aspartate, indicating that Cl⁻ is the primary permeant ion (Fig. 6F).

CG10353-mCherry fusion protein was concentrated at the plasma membrane of HEK293T cells, as shown by co-localization with FITC-WGA, which specifically binds to GlcNAc- and sialic acid-containing glycoconjugates displayed on the extracellular surface of mammalian cells. In contrast, we did not detect such co-localization for DmTMEM16O-mCherry or mCherry alone (Fig. 6G). These results suggest that DmTMEM16O may require specific *D. melanogaster* ORN proteins absent in the heterologous system for efficient localization and function at the plasma membrane.

## Discussion

In this study, we identify DmTMEM16O as a key regulator of ORN response dynamics in *D. melanogaster*. DmTMEM16O is selectively enriched in Orco-positive ORNs in both larval and adult olfactory organs, and its loss leads to a pronounced defect in response termination. Across multiple sensillum types, *DmTMEM16O* mutants exhibit prolonged odor-evoked spiking and slow receptor potential decay after stimulus offset, resulting in persistent neuronal activity, impaired temporal resolution during repeated stimulation, and reduced behavioral responses to both attractive and aversive odorants. These findings establish DmTMEM16O as a critical determinant of the kinetics, rather than the amplitude, of ORN responses.

A central insight from our data is that DmTMEM16O regulates the membrane time constant of ORNs by contributing to total membrane conductance during the decay phase of the response. The prolonged decay times observed in mutants, together with largely preserved spiking frequency, indicate that DmTMEM16O primarily accelerates repolarization after odor offset. The most parsimonious model is that DmTMEM16O contributes a repolarizing conductance that is engaged during or immediately after odor stimulation. Given the low intracellular Cl⁻ concentration in insect ORNs, about 24 mM in moths [23] and 20 mM in flies [24], activation of a chloride conductance would drive Cl⁻ influx, promoting membrane repolarization. This is consistent with two previous studies on whole cell patch-clamp recording of odor-activated moth ORNs [25] and *in vivo* imaging of intracellular Cl⁻ with odor-activated *Drosophila* ORNs [24]. In this context, DmTMEM16O would function to increase total membrane conductance and accelerate decay kinetics, thereby sharpening the temporal profile of ORN responses (Fig. 7A). Since insect sensillum lymph contains unusually high K⁺ concentrations [26], odor-evoked depolarization of ORNs likely involves mixed cation influx through OR-Orco channels under ionic conditions distinct from those of conventional neurons, potentially increasing the importance of active repolarization mechanisms. This model explains not only the prolonged decay but also the failure of mutant neurons to resolve repeated odor stimuli, as a longer membrane time constant leads to temporal summation and loss of stimulus separation.

**Figure 7.**
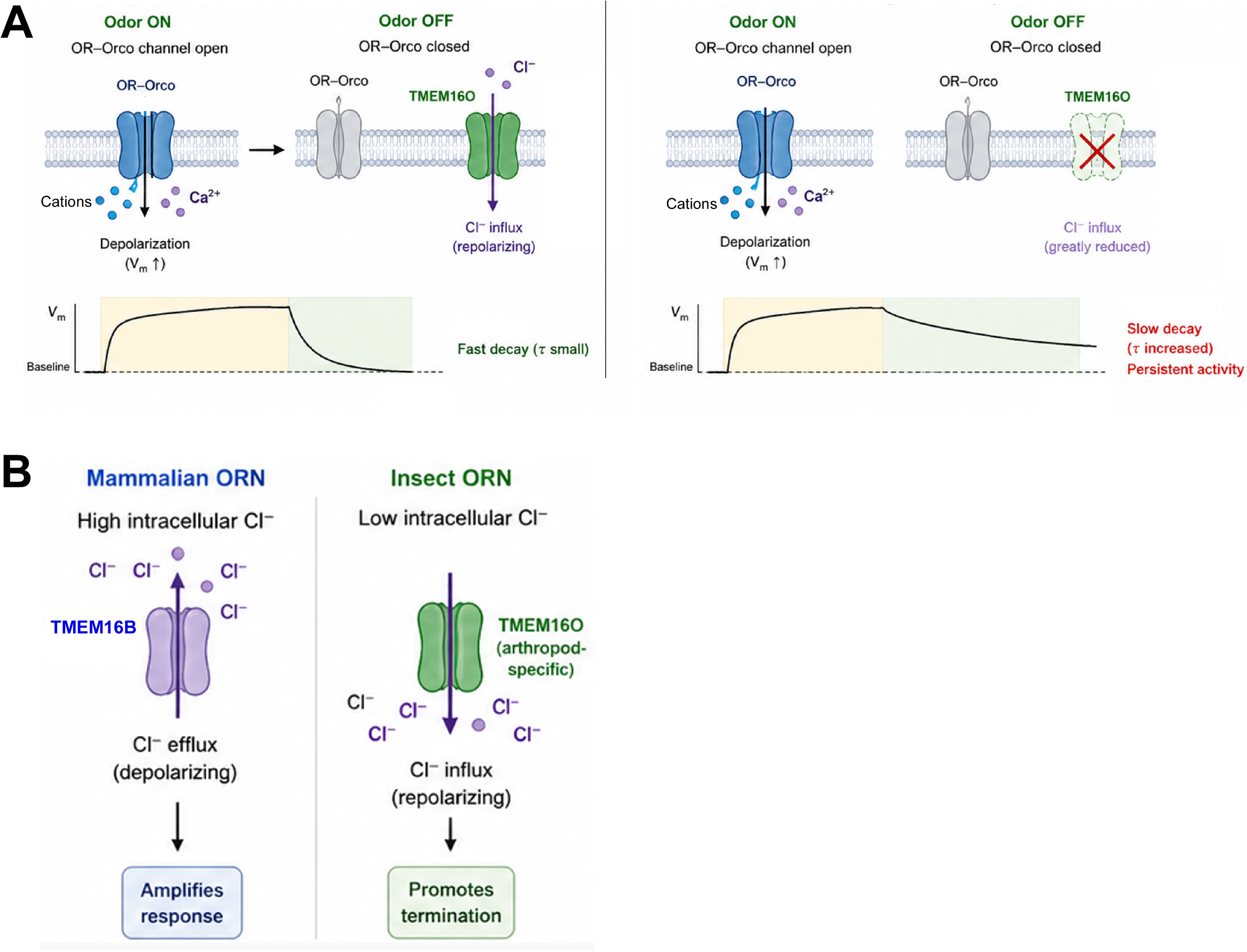
TMEM16O accelerates termination of olfactory responses by increasing membrane conductance in insect ORNs. (A) In WT ORNs, odor stimulation activates OR–Orco channels, leading to cations and Ca^2+^ influx and membrane depolarization. Upon odor offset, TMEM16O-mediated Cl⁻ influx contributes to rapid repolarization, resulting in fast decay of receptor potentials. In DmTMEM16O mutants, activation is preserved, but loss of TMEM16O reduces membrane conductance during odor offset, leading to prolonged depolarization and slower decay kinetics. (B) Chloride currents play opposite roles in mammalian and insect ORNs. In mammals, TMEM16B mediates Cl⁻ efflux due to high intracellular Cl⁻, amplifying depolarization. In insects, TMEM16O is proposed to mediate Cl⁻ influx, promoting repolarization.

This proposed function contrasts strikingly with the role of TMEM16B in mammalian ORNs. TMEM16B opening results in Cl⁻ efflux, which further depolarizes the neuron and amplifies the odor response. Thus, TMEM16B plays roles for amplification of the odorant response [12], modulation of ORN firing [27], and response kinetics and adaptation [28]. TMEM16B-deficient mouse ORNs display faster response termination, allowing them to fire action potentials more reliably during rapid repeated stimulation [29]. This phenotype is indeed opposite from DmTMEM16O mutant ORNs. DmTMEM16O therefore likely plays a role opposite to that of TMEM16B, acting to terminate rather than amplify sensory responses (Fig. 7B). This difference highlights how similar channel families can be repurposed to serve distinct physiological roles depending on ionic gradients and signaling architecture.

Although DmTMEM16O belongs to the TMEM16 family, our heterologous expression experiments indicate that it does not function as a conventional CaCC under standard conditions. Unlike the *Drosophila* TMEM16A/B homolog CG10353, DmTMEM16O did not generate detectable Ca^2+^-dependent currents in HEK293T cells and failed to localize efficiently to the plasma membrane. These results suggest that DmTMEM16O either requires ORN-specific factors for proper trafficking and activation or fulfills a non-canonical role within the TMEM16 family. One possibility is that DmTMEM16O operates within specialized microdomains near OR–Orco complexes, where local Ca^2+^ dynamics or protein interactions differ substantially from those in heterologous systems. Alternatively, DmTMEM16O may function as a lipid scramblase or regulatory protein that indirectly modulates membrane conductance rather than forming an ion channel itself. Based on Fly Cell Atlas data, CG10353 is highly expressed in midgut associated cells and Malpighian tubule cells, suggesting that it is unlikely to function as CaCC in ORN.

The defects in temporal coding observed in *DmTMEM16O* mutants provide an important link between cellular physiology and behavior. In natural environments, odor signals are highly intermittent, and accurate perception depends on the ability of ORNs to resolve rapid fluctuations. In WT flies, fast repolarization allows neurons to reset between odor encounters, preserving temporal fidelity. In mutants, prolonged depolarization leads to response overlap and reduced contrast between stimulus onset and offset. These changes likely degrade the quality of sensory information transmitted to higher-order circuits, explaining the reduced behavioral responses despite sustained neuronal activity. Similarly, pheromone-binding protein–deficient male silk moths exhibit reduced temporal resolution in their antennal responses and require significantly longer to locate pheromone sources compared to wild-type males [30].

The evolutionary restriction of TMEM16O to arthropods suggests that it represents a lineage-specific adaptation to the unique properties of insect olfactory signaling. Unlike vertebrates, which rely on GPCR-based transduction, insects use ionotropic OR–Orco channels that produce rapid depolarization. This mode of signaling likely imposes distinct constraints on response termination, necessitating mechanisms that can quickly restore membrane potential. TMEM16O appears well suited to this role, providing a conductance that accelerates repolarization and ensures high temporal resolution. This function represents a mechanistically and evolutionarily distinct solution to sensory signal termination in insects. In addition to ORNs in the DOG, DmTMEM16O was also detected in larval pharyngeal sensory neurons. This is consistent with Fly Cell Atlas data that DmTMEM16O is also expressed in labellar gustatory receptor neurons (GRNs) and the mechanosensory neurons of Johnston’s organ, suggesting that its function is not restricted to olfaction. Labellar GRNs require rapid adaptation during feeding, whereas Johnston’s organ neurons encode high-frequency mechanical stimuli such as sound and antennal movement. In both systems, delayed response termination would impair temporal fidelity. These observations raise the possibility that TMEM16O functions more broadly as an arthropod-specific regulator that accelerates recovery of activated peripheral sensory neurons.

## Materials and Methods

### Construction of plasmids and generation of transgenic flies

To generate the DmTMEM16O-Gal4 construct, the 5′ upstream region of DmTMEM16O was PCR-amplified from Oregon-R genomic DNA using the primers DmTMEM16O promoter-5-BamHI and DmTMEM16O promoter-3-HindIII. The PCR product was digested with BamHI and HindIII and subcloned into BglII/HindIII-digested pGAL4attB vector [31]. The resulting construct was integrated into the attP40 landing site on the second chromosome (25C6) using phiC31 integrase-mediated transformation [32].

For IR25a-tdTomato, the 5′ upstream region of IR25a was amplified from Oregon-R genomic DNA using IR25a promoter-5 and IR25a promoter-3 primers. The amplified fragment was phosphorylated and subcloned into HindIII/BglII-digested pJFRC2 vector [33] (Addgene #26214), in which mCD8-GFP had been replaced with tdTomato. HindIII and BglII sites were blunt-ended using Klenow fragment prior to ligation. The resulting construct was integrated into the attP2 landing site on the third chromosome (68A4) using phiC31 integrase.

To generate UAS-DmTMEM16O, the DNA fragment encoding the DmTMEM16O open reading frame (ORF) was excised from a mammalian expression vector using EcoRI and XbaI and subcloned into the corresponding sites of pUASTB [34] (Addgene #18944). The construct was integrated into the attP40 landing site on the second chromosome by phiC31 integrase.

### Generation of DmTMEM16O-deficient flies by CRISPR

DmTMEM16O mutant flies were generated using CRISPR/Cas9 as described previously [35]. Guide RNAs (gRNAs) targeting exon 3 and exon 10 were generated by annealing complementary oligonucleotides (DmTMEM16O-exon3-gRNA-F/R and DmTMEM16O-exon10-gRNA-F/R) and subcloning them into BbsI-digested pBFv-U6.2 and pBFv-U6.2B vectors, respectively. The EcoRI/NotI fragment containing the exon 3 gRNA cassette was subsequently inserted into pBFv-U6.2B carrying the exon 10 gRNA cassette to generate a dual-gRNA expression construct.

The plasmid expressing both gRNAs was injected into *y^1^ v^1^ nos-phiC31; attP40* embryos [32]. Surviving G0 males were individually crossed to *y^2^ cho^2^ v^1^* virgin females, and transformants were balanced using *Sp/CyO*. Females carrying the *U6-DmTMEM16O gRNA* construct were then crossed with nos-Cas9 males (*y^2^ cho^2^ v^1^; attP40[nos-Cas9]/CyO*). Founder males carrying both the gRNA construct and nos-Cas9 were crossed with *w; Dr/TM6B* virgin females.

Deletion of the genomic region between exon 3 and exon 10 was screened by wing PCR using primers DmTMEM16O-388F and DmTMEM16O-3094R. Flies carrying the deletion allele were identified by the presence of a smaller PCR product in addition to the wild-type *TM6B*-derived band. Homozygous mutant stocks were established following appropriate crosses, and deletion breakpoints in each stock were determined by sequencing PCR amplicons generated using the same primer pair. The largest deletion allele, *DmTMEM16O^Δ153–1016^*, was backcrossed to Oregon-R for five generations to isogenize the genetic background.

### Drosophila genetics

*DmTMEM16O-Gal4* flies were crossed with *y^1^ w; 20XUAS-6XGFP VK00018 (53B2)/CyO* (Bloomington stock #52261). *DmTMEM16O-Gal4/20XUAS-6XGFP* females were then crossed with *w; wg^Sp-1^/CyO; MKRS/TM2* males. Recombinant flies carrying *DmTMEM16O-Gal4* and *20XUAS-6XGFP* on the second chromosome were selected and established.

*DmTMEM16O-Gal4 20XUAS-6XGFP/CyO; TM2/TM6B* females were crossed with *y^1^ w; wg^Sp-1^/CyO; 20XUAS-6XGFP attP2 (68A4)* males (Bloomington stock #52262). *DmTMEM16O-Gal4 20XUAS-6XGFP/CyO; 20XUAS-6XGFP/TM6B* males and females were then crossed to establish the *DmTMEM16O-Gal4 20XUAS-6XGFP; 20XUAS-6XGFP* stock.

To generate flies expressing IR25a-tdTomato, *DmTMEM16O-Gal4 20XUAS-6XGFP/CyO; TM2/TM6B* females were crossed with *w; wg^Sp-1^/CyO; IR25a-tdTomato* males. *DmTMEM16O-Gal4 20XUAS-6XGFP/CyO; IR25a-tdTomato/TM6B* males and females were then crossed to establish *DmTMEM16O-Gal4 20XUAS-6XGFP; IR25a-tdTomato* flies.

To generate flies expressing Orco-RFP, *DmTMEM16O-Gal4 20XUAS-6XGFP/CyO; TM2/TM6B* females were crossed with *w; UAS-mCD8::GFP/CyO; Orco-RFP* flies (Bloomington stock #63045). *DmTMEM16O-Gal4 20XUAS-6XGFP/CyO; Orco-RFP/TM6B* males and females were subsequently crossed to establish *DmTMEM16O-Gal4 20XUAS-6XGFP; Orco-RFP* flies.

For rescue experiments, males of *w; UAS-DmTMEM16O/CyO; TM2/TM6B* or *DmTMEM16O-Gal4/CyO; TM2/TM6B* were crossed with *w, wg^Sp-1^/CyO; DmTMEM16O^Δ153–1016^/TM6B* females. *UAS-DmTMEM16O/CyO; DmTMEM16O^Δ153– 1016^/TM6B* males and females were crossed to establish *UAS-DmTMEM16O; DmTMEM16O^Δ153–1016^* flies. Similarly, *DmTMEM16O-Gal4/CyO; DmTMEM16O^Δ153– 1016^/TM6B* males and females were crossed to establish *DmTMEM16O-Gal4; DmTMEM16O^Δ153–1016^* flies.

### Confocal live imaging

For *in vivo* imaging of first instar larvae, larvae were immobilized by anesthetization with neat ethyl acetate as described previously [36]. Briefly, selected first instar larvae were gently washed with water to remove residual food and dried on Kimwipes. Washed larvae were transferred into a larval cage, which was placed inside an anesthetic chamber containing cotton wool soaked with 15 mL ethyl acetate for 88–95 s. Anesthetized larvae were mounted in phosphate-buffered saline (PBS) between a coverslip and microscope slide, with two additional coverslips positioned on either side of the larvae to prevent compression. For adult imaging, dissected fly heads were mounted ventral side up in PBS between two coverslips, with a third coverslip placed on top to avoid flattening.

Images were acquired using a Zeiss LSM880 laser-scanning confocal microscope. For larval co-localization experiments, Z-stacks were obtained to identify the brightest focal planes for each fluorophore. Raw confocal images were processed using Fiji (cropping, channel merging, brightness/contrast adjustment, and maximum intensity projection of Z-stacks) and Adobe Photoshop (overlay of brightfield and fluorescence images).

### Single sensillum recording

Odorants were obtained from Sigma-Aldrich (UK) and were of the highest purity available; chiral compounds were racemic mixtures. For antennal basiconic sensilla recordings, odorant concentrations were selected to elicit approximately half-maximal responses (EC50) in individual ORNs. For maxillary palp recordings, odorants were diluted to 10⁻² (v/v) from pure in paraffin oil, as described previously [37]. Odor cartridges were prepared by applying 30 μL diluted odorant solution onto a 15-mm filter paper disc inserted into a 150-mm glass Pasteur pipette (2 mL volume). The cartridge was capped with a 1000 μL pipette tip, and both ends were sealed with Parafilm until use. A tube carried an air stream directed at the fly, and odorants were delivered by injecting air through the cartridge placed with its tip into an aperture of the tube so the air stream carried the volatile compounds to the fly. Each odor cartridge was used for a maximum of three stimulations [22].

Adult flies (2–10 days old) were immobilized in 100 μm pipette tips, and the antenna or maxillary palp was stabilized against a coverslip using the tip of a glass capillary [38]. Extracellular recordings from ORNs were obtained by inserting a recording electrode into the sensillum lymph surrounding the dendrites, while a reference electrode was inserted into the eye [22]. Both unfiltered traces and 500 Hz high-pass filtered traces were recorded.

The antennae and maxillary palps were visualized at 1000× magnification using an Olympus BX51WI upright microscope (Olympus, UK), allowing individual sensilla to be clearly identified. Recording electrodes were fabricated from glass capillaries using a micropipette puller and drawn to <1 μm tip diameter. Electrodes were filled with sensillum lymph Ringer solution [39] and connected to Ag/AgCl silver wire electrodes. No more than three sensilla were recorded from a single fly. Each sensillum was stimulated with multiple odorants with at least 15 s between presentations.

### Calculation of spikes/s and decay time constant (τoff)

Signals were recorded beginning 1 s before odor stimulation. Spike numbers during the 1 s pre-stimulus period and the 0.5 s stimulation period were quantified offline using custom MATLAB scripts. Spike rate (spikes/s) of individual neurons was calculated by counting spikes within the 0.5 s stimulation window, multiplying by 2, and subtracting the number of spikes detected during the 1 s pre-stimulus period. For repeated-stimulation experiments, flies were exposed to 5 cycles of 100 ms odor stimulation separated by 100 ms rest intervals, and spikes during each ON and OFF phase were quantified separately. The decay time constant (τoff) was used to quantify the rate of response termination following odor offset. τoff represents the time required for the firing rate to decay to 37 % (1/e) of its peak value [40]. Spike detection was performed on high-pass filtered traces using the MATLAB *findpeaks* function with a minimum inter-spike interval of 5 ms to avoid double counting. Detection thresholds were determined individually for each recording based on pre-stimulus noise levels. Noise standard deviation (σ_noise_) was calculated from the baseline interval (t = 0–0.9 s; odor onset at t = 1 s), and threshold sensitivity was tested over a range of 0.5×–10×σ_noise_. The final threshold was selected as the lowest value that reduced pre-stimulus spike detection to near zero while preserving maximal odor-evoked spike counts. Because SSR recordings consist of discrete spike trains, instantaneous firing rates were estimated from single-trial spike times using Gaussian kernel density estimation [41]:

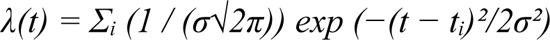

where *tᵢ* is the time of the *i*-th spike and σ is the kernel width. Kernel width was initially set to one-fifth of the decay analysis window and adjusted within the range of one-eighth to one-fourth of the window duration according to spike count. Wider kernels were used for recordings with fewer than 10 spikes, whereas narrower kernels were used for recordings with 20 or more spikes. Kernel widths were constrained between 50 and 500 ms, and final values were confirmed by visual inspection to ensure smooth decay curves without over-smoothing. For each sensillum–odorant combination, all recordings were analyzed using identical parameters across genotypes, including kernel width, decay window duration, and *τ_off_* fitting bounds. Baseline firing was corrected by subtracting the mean pre-stimulus kernel density (t = 0–0.9 s), and negative values were set to zero. Following odor offset, the decay of ORN firing rates was fitted with a single exponential function:

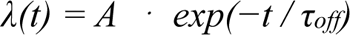

where *A* is the peak firing rate at odor offset, *t* is the elapsed time after odor offset (defined as t = 0 at odor offset, corresponding to 1.5 s after trial onset), and *τ_off_* is the decay time constant. The model was fitted to baseline-corrected kernel density estimates using nonlinear least-squares fitting (MATLAB *fit* function). Fitting was performed over the decay analysis window from the peak response to T_REST_. Lower and upper bounds for *τ_off_* were set to 0.01 s and T_REST_, respectively, and applied consistently across genotypes for each sensillum–odorant combination. Goodness of fit was evaluated using the coefficient of determination (R²), and recordings with R² > 0.80 were included in the analysis.

### T-maze assay

T-maze assays were performed with modifications from previous studies [42, 43]. Mated female and male flies aged 3–4 days (n = 10–15 per test) were starved for 24 h with access to water. On the following day, the starved flies (4–5 days old) were anesthetized on ice and introduced into the starting chamber of a T-shaped glass tube consisting of a 5.0 cm starting chamber and two 4.5 cm arms with an inner diameter of 5.0 mm. Each arm was connected through a silicone tube (1.8 cm length, 8.0 mm diameter) to a 2 mL glass vial positioned at the end of the apparatus. To prevent flies from directly contacting (i.e. tasting) the odorant or solvent solution while still allowing volatile diffusion into the T-maze arm, a small nylon mesh filter was placed between each arm and the connected glass vial. One vial contained 5 μL diluted odorant solution at either 10^-2^or 10^-4^ (v/v) dilution [42, 44], whereas the opposite vial contained 5 μL solvent control (paraffin oil). Solutions were applied onto 6.0 × 6.0 mm filter paper pieces placed inside the vials.

After introducing anesthetized flies into the starting chamber, the bottom opening of the T-maze was sealed with Parafilm to prevent escape. Flies typically resumed movement within 3–4 min, and this recovery period was excluded from the assay duration. Once recovered, the T-maze apparatus was positioned vertically. The numbers of flies in the arm connected to the odorant-containing vial (“odorant side”), the arm connected to the solvent control vial (“solvent side”), and the starting chamber were counted after 10 min. Flies remaining in the starting chamber were excluded from the analysis. Olfactory preference was quantified using the response index (RI):

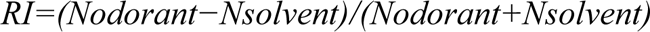

where *Nodorant* and *Nsolvent* represent the numbers of flies on the odorant and solvent sides, respectively. During experiments, T-maze tubes were placed on a white surface under white light illumination. All flies were discarded after testing.

### Plasmid DNA constructs for expression of DmTMEM16O, CG10353, and mCherry fusion proteins in HEK293T cells

An NsiI–MluI fragment from pCMVTag5A containing the CMV promoter, multicloning site, and SV40 polyadenylation sequence was first subcloned into the corresponding sites of pGEM-T Vector. The DNA fragment encoding mCherry was PCR amplified using the primers mCherry-5-ClaI and mCherry-3-XhoI with mCherry-Farnesyl-5 (a gift from Michael Davidson (Addgene plasmid # 55045; http://n2t.net/addgene:55045; RRID:Addgene_55045) as a template, and subcloned into the ClaI and XhoI sites of the above vector.

The full-length DmTMEM16O open reading frame (ORF) was obtained by nested PCR. The first PCR was performed using the outer primers DmTMEM16O-5-outer and DmTMEM16O-3-outer with reverse-transcribed RNA from adult *D. melanogaster* heads as a template. The second PCR used DmTMEM16O-5-EcoRI and DmTMEM16O-3-ClaI primers with the first PCR product as a template. The resulting amplicon was subcloned into the EcoRI and ClaI sites of the mCherry vector to generate DmTMEM16O-mCherry. The full-length CG10353 ORF was amplified using CG10353-5-NotI and CG10353-3-HindIII primers with reverse-transcribed RNA from *D. melanogaster* embryos as a template. The PCR product was subcloned into the NotI and HindIII sites of the same vector to generate CG10353-mCherry.

To generate untagged CG10353, PCR was performed using primers CG10353-1448F and CG10353-3-Stop-HindIII with the CG10353-mCherry plasmid as a template. The amplicon was digested with BamHI and HindIII and subcloned into the same sites of the mCherry vector after introduction of a stop codon at the end of the CG10353 ORF. To generate untagged DmTMEM16O, PCR was performed using primers DmTMEM16O-5-EcoRI and DmTMEM16O-3-Stop-XbaI with the DmTMEM16O-mCherry plasmid as a template. The resulting fragment was digested with EcoRI and XbaI and subcloned into pAc5.1/V5-His B, in which the *Drosophila actin 5C* promoter had been replaced with the CMV promoter [45]. All primers used in this study are listed in Supplementary Table 1.

### FITC-WGA staining and immunofluorescence

HEK293T cells transiently expressing mCherry, DmTMEM16O-mCherry, or CG10353-mCherry were cultured in 8-well chamber slides. Cells were washed with ice-cold PBS and incubated with 5 μg/mL FITC-conjugated wheat germ agglutinin (FITC-WGA) in PBS for 15 min on ice to label the plasma membrane. Following three washes with PBS, cells were fixed with 4% paraformaldehyde for 15 min at room temperature and permeabilized with 0.1% Triton X-100 for 5 min. Cells were blocked with 5% BSA in PBS for 30 min and incubated overnight at 4°C with rabbit anti-mCherry antibody (Proteintech). After three PBS washes, cells were incubated with Alexa Fluor 594-conjugated anti-rabbit IgG secondary antibody (Thermo Fisher) for 3 h at room temperature. Cells were then washed three times with PBS, mounted, and imaged using confocal microscopy.

### Electrophysiology

HEK293T cells were maintained in DMEM supplemented with 10% fetal bovine serum and penicillin–streptomycin at 37°C in 5% CO₂. HEK293T cells were transiently transfected with 1 μg plasmid DNA encoding DmTMEM16O, CG10353, or a control construct together with 0.1 μg pGreen Lantern-1 vector using Lipofectamine and Plus Reagent (Thermo Fisher). Whole-cell patch-clamp recordings were performed 24 h after transfection at room temperature.

Recording pipettes (3–5 MΩ) were filled with intracellular solution containing 140 mM NaCl (Fig. 6B), NMDG-Cl (Fig. 6A, C, and E), or NMDG-aspartate (Fig. 6D), together with 10 mM HEPES and 5 mM EGTA. Free Ca²⁺ concentration was adjusted to 100 μM for experiments shown in Fig. 6A–D and F, whereas Ca²⁺ was omitted in Ca²⁺-free recordings shown in Fig. 6E. The pH was adjusted to 7.3.

The extracellular bath solution contained the same ionic composition as the corresponding intracellular pipette solution. Cells were voltage-clamped at 0 mV, and voltage step pulses from −100 to +100 mV in 20 mV increments were applied for 100–500 ms at 3 s intervals. Currents were filtered at 1–2 kHz, digitized, and analyzed using standard electrophysiological software. Current amplitudes and current–voltage (I–V) relationships were subsequently quantified.

## CRediT authorship contribution statement

Tatsuhiko Kadowaki conceived and designed research strategy and wrote the paper. Ying Lei, Jing Lei, and Tianbang Li performed the experiments. Makoto Tominaga and Wynand Van Der Goes Van Naters supervised the experiments. Wynand Van Der Goes Van Naters edited the manuscript.

## Acknowledgements

This work was supported by Jinji Lake Double Hundred Talents Programme to TK. The funder had no role in study design, data collection and analysis, decision to publish, or preparation of the manuscript.

## Ethics statement

Not applicable

## Availability of data and materials

All data is included in the manuscript and Supplementary material.

---

**Supplementary Figure 1.** Deleted region of DmTMEM16O in the *DmTMEM16O^Δ153–1016^* mutant DmTMEM16O is a 1229 amino-acid protein containing an Anoctamin dimerization domain (Anoct_dimer; amino acids 340–573) and an Anoctamin domain with ten transmembrane segments (amino acids 576–1149). The deleted region in *DmTMEM16O^Δ153–1016^*(amino acids 153–1016) includes the entire Anoctamin dimerization domain and transmembrane segments 1–9 of the Anoctamin domain.

**Supplementary Figure 2.** Spiking frequencies of ORNs in ab1, ab2, ab3, and pb1 sensilla of WT and *DmTMEM16O^Δ153–1016^* flies. Spiking frequencies of individual ORNs in ab1, ab2, ab3, and pb1 sensilla in response to methyl salicylate (A), ethyl acetate (B), ethyl 3-hydroxybutyrate (C), ethyl butyrate (D), 2-heptanone (E), racemic 3-octanol (F), and 4-methyl phenol (G) were comparable between WT and *DmTMEM16O^Δ153–1016^* flies (ns, Brunner–Munzel test).

**Supplementary Table 1.** List of primers used in this study

