## Supplementary figures and images for "An arthropod-specific TMEM16 protein accelerates olfactory response termination in *Drosophila*"

### Supplementary Figure 1.pdf

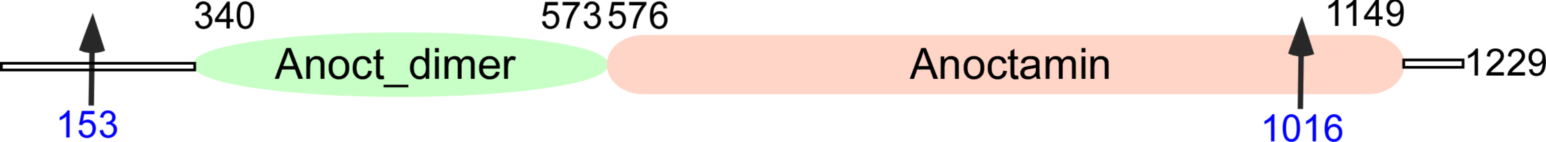

### Supplementary Figure 2.pdf

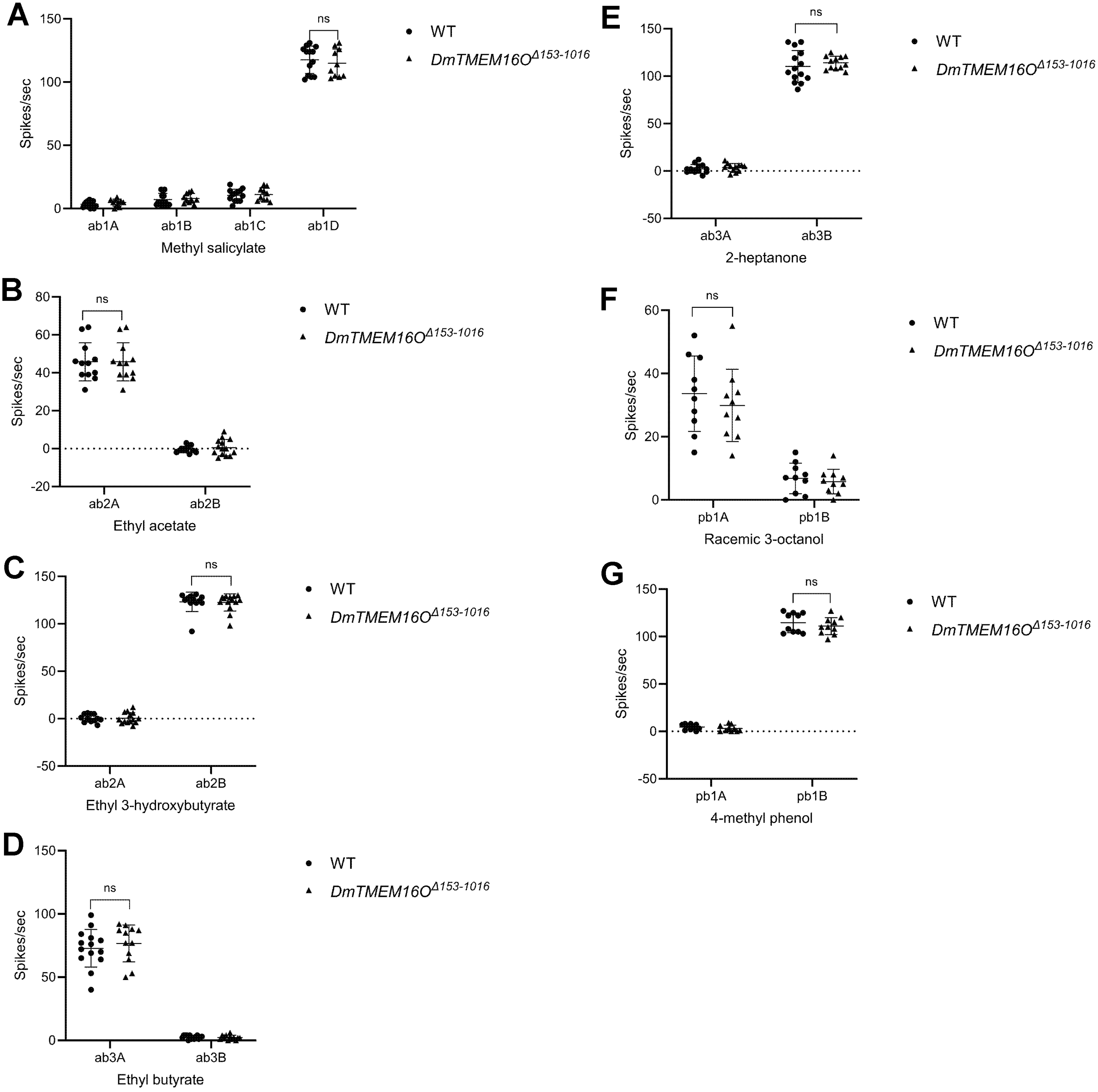
